# Direct piriform-to-auditory cortical projections shape auditory-olfactory integration

**DOI:** 10.1101/2024.07.11.602976

**Authors:** Nathan W. Vogler, Ruoyi Chen, Alister Virkler, Violet Y. Tu, Jay A. Gottfried, Maria N. Geffen

**Author notes:** Corresponding Author 3450 Hamilton Walk, Stemmler Hall G10, Philadelphia, PA 19104.

## Abstract

In a real-world environment, the brain must integrate information from multiple sensory modalities, including the auditory and olfactory systems. However, little is known about the neuronal circuits governing how odors influence and modulate sound processing. Here, we investigated the mechanisms underlying auditory-olfactory integration using anatomical, electrophysiological, and optogenetic approaches, focusing on the auditory cortex as a key locus for cross-modal integration. First, retrograde and anterograde viral tracing strategies revealed a direct projection from the piriform cortex to the auditory cortex. Next, using *in vivo* electrophysiological recordings of neuronal activity in the auditory cortex of awake male or female mice, we found that odors modulate auditory cortical responses to sound. Finally, we used *in vivo* optogenetic manipulations during electrophysiology to demonstrate that olfactory modulation in auditory cortex, specifically, odor-driven enhancement of sound responses, depends on direct input from the piriform cortex. Together, our results identify a novel role of piriform-to-auditory cortical circuitry in shaping olfactory modulation in the auditory cortex, shedding new light on the neuronal mechanisms underlying auditory-olfactory integration.

**Significance Statement:** All living organisms exist within multisensory environments, yet there is a lack in our understanding of how the brain integrates multisensory information. This work highlights a direct piriform-to-auditory cortical circuit governing auditory-olfactory integration. Our results shed new light on a relatively understudied area of multisensory research, promising a more robust understanding of how animals and humans perceive and interact within complex environments.

## Introduction

A fundamental role of the central nervous system is to integrate information across multiple sensory modalities to optimize decision-making and behavior. Indeed, rarely does any individual sensory modality operate alone and unabetted: humans and animals must integrate combinations of sounds, smells, visual input, tactile and proprioceptive information (Gottfried and Dolan, 2003; Deroy et al., 2016; Angeloni and Geffen, 2018; Choi et al., 2018; Wallace et al., 2020). When combining sounds and smells, the brain must integrate information from both the auditory and olfactory systems. Auditory-olfactory integration is an ethologically important domain of multisensory integration. When a cat approaches a mouse, the mouse may use the combination of sounds and smell of the cat to identify it as a predator (Halene et al., 2009). Moreover, combinations of auditory and olfactory stimuli are crucial for pup retrieval behavior in mice, and play critical roles during development — bonds between parents and offspring form based on body odors and vocalization sounds (Marlin and Froemke, 2017; Tasaka et al., 2020; McRae et al., 2023). In humans, the sounds of eating food, such as crunchy potato chips, affect perception of the pleasantness of the food odor, such as potato chip odor/staleness (Seo and Hummel, 2011; Seo et al., 2014). However, the circuits and mechanisms underlying auditory-olfactory integration have remained elusive.

Previous studies examining interactions between audition and olfaction demonstrated cross-modal activation of piriform and auditory cortical areas (Smith et al., 2009; Wesson and Wilson, 2010; Cohen et al., 2011; Varga and Wesson, 2013; Cohen and Mizrahi, 2015; Gnaedinger et al., 2019; Olofsson et al., 2019; Tasaka et al., 2020; Gilday and Mizrahi, 2023; Wu et al., 2023). In the auditory cortex (ACx) of lactating mother mice, pup odors reduce spontaneous activity and increase evoked activity in response to pup ultrasonic vocalizations (Cohen et al., 2011; Cohen and Mizrahi, 2015). Furthermore, learned associations between an odor and a subsequent sound modulate single-neuron responses in the auditory cortex (Gilday and Mizrahi, 2023). Together, these studies suggest that the ACx serves as a locus for auditory-olfactory cross-modal integration, specifically regarding the olfactory modulation of sound processing.

While these studies point to an involvement of the ACx in auditory-olfactory integration, much remains unknown regarding where and how these cross-modal inputs converge. By comparison, previous research on audio-visual integration has found that concurrent visual stimuli improve sound detection and encoding in ACx, facilitated by cross-cortical connections between visual cortex and ACx (Bizley et al., 2016; Atilgan et al., 2018; Meijer et al., 2018; Zhou et al., 2020; Han et al., 2021). Moreover, audio-visual integration crucially depends on the temporal binding of the combined audio-visual stimuli (Meijer et al., 2017, 2019). Similar studies on auditory-olfactory integration have been limited by experimental challenges in delivering combinations of odors and sounds with temporal precision, as well as a lack of knowledge of the brain circuitry underlying auditory-olfactory interactions. Notably, reports of potential cross-modal anatomical connections linking the piriform cortex (Pir) and ACx have been inconsistent. Some have found that neurons in Pir display sound-evoked responses (Varga and Wesson, 2013), and that ACx projects to Pir but not vice versa (Budinger et al., 2006; Budinger and Scheich, 2009). Others have reported a sparse but detectable direct projection from Pir to ACx (Costa et al., 2017; Tasaka et al., 2020), yet its potential role in odor-driven modulation of ACx is unclear. Identifying and characterizing these pathways are important first steps in deciphering how the brain integrates auditory and olfactory stimuli.

Here, we confirm the direct Pir-to-ACx projection using multiple circuit tracing strategies and examine its functional role in auditory-olfactory integration in ACx. We explore the impact of this pathway on olfactory modulation of auditory processing, using a novel experimental setup for delivering odors and sounds with temporal precision to awake mice during *in vivo* electrophysiological recordings in ACx. Our findings reveal that the Pir-to-ACx pathway contributes specifically to odor-mediated enhancement of sound responses in ACx.

## Materials and Methods

### Mice

For anatomical circuit tracing experiments, we used 5 male C57BL/6J mice (The Jackson Laboratory; strain #000664; RRID:IMSR_JAX:000664; aged 8-16 weeks), 2 male Cdh23 mice (B6N(Cg)-Cdh23^tm2.1Kjn^/Kjn; The Jackson Laboratory, strain #:018399; RRID:IMSR_JAX:018399; aged 22 weeks), and 13 Ai14 Cre reporter mice (progeny of Cdh23 mice crossed with Ai14 mice [B6.Cg-Gt(ROSA)^26Sortm14(CAG-tdTomato)Hze^/J; The Jackson Laboratory, strain #:007914; RRID:IMSR_JAX:007914], 7 male, 6 female, aged 18-26 weeks). For electrophysiological recordings, we used 18 mice: 4 C57BL/6J mice (2 male, 2 female; aged 12-17 weeks at time of recording); and 14 Cdh23 mice (5 male, 9 female; aged 12-26 weeks). Cdh23 mice were used for most experiments because they do not exhibit age-related hearing loss which is common in C57BL/6J mice (Johnson et al., 2017). Any C57BL/6J mice were used by 17 weeks of age. Mice were housed at 28 °C with a 12-h reversed light cycle and food and water provided *ad libitum*. Experiments were performed during the animals’ dark cycle. All experimental procedures were approved by the Institutional Care and Use Committee at the University of Pennsylvania and in accordance with National Institutes of Health guidelines.

### Stereotaxic surgery and virus injections

Mice were anesthetized under isoflurane (1-3%). Prior to surgery, mice were administered subcutaneous doses of buprenorphine (0.05-0.1 mg/kg Buprenex or 1.3 mg/kg Ethiqa), dexamethasone (0.2 mg/kg), and bupivicane (local; 2 mg/kg). A small craniotomy (∼1 mm diameter) was made above the respective ACx or Pir injection sites. To label ACx-projecting neurons in Pir in Cdh23 mice, we injected AAVrg-hSyn-eGFP (Addgene #50465-AAVrg; RRID:Addgene_50465; 1.7x10^13^ GC/mL; 250 nL) into left ACx (4.3 mm ML and -2.65 mm AP [relative to Bregma], -0.7 mm DV [relative to brain surface]). To label ACx-projecting neurons in Pir in Ai14 mice, we injected AAVrg-hSyn-Cre-WPRE-hGH (Addgene #105553-AAVrg; RRID:Addgene_105553; 1.9 x10^13^ GC/mL; 150-250 nL) into left ACx. To label Pir-recipient neurons in ACx in Ai14 mice, we injected AAV1-hSyn-Cre-WPRE-hGH (Addgene #105553-AAV1; RRID:Addgene_105553; 1.9 x10^13^ GC/mL; 150-180 nL) into left Pir (3.9 mm ML, -1.5 mm AP, - 4.00 DV). Viruses were delivered through 30-40 μm diameter glass micropipettes connected to a syringe pump (Harvard Apparatus Pump 11 Elite; 60 nL/min infusion rate). For optogenetic experiments, in Cdh23 or C57BL/6J mice we injected AAVrg-hSyn-Cre-WPRE-hGH (300 nL) into left ACx and AAV5-hSyn1-SIO-stGtACR1-FusionRed (Addgene #105678-AAV5; RRID:Addgene_105678; 2.1 x10^13^ GC/mL; 300 nL) or AAV5-FLEX-tdTomato (Addgene #28306-AAV5; RRID:Addgene_28306; 2.1 x10^13^ GC/mL; 300 nL) into left Pir (3.9 mm ML, -1.0 mm AP, - 4.00 mm DV). A fiber optic cannula (ThorLabs CFML52L05; Ø200 µm Core, 0.50 NA, L=5mm) was implanted over left Pir (3.9 mm ML, -1.0 mm AP, -3.6 mm DV) and secured with dental cement (C&B Metabond). For all electrophysiological recordings, a ground pin was inserted via small craniotomy over the left frontal cortex, and a custom-made stainless steel headplate (eMachine Shop) was secured to the skull using dental cement and acrylic (Lang Dental). After all surgeries, we subcutaneously administered meloxicam (5 mg/kg) and an antibiotic (Baytril 5 mg/kg) for 4 days during postoperative care. After virus injections, to allow adequate time for virus expression, we waited at least two weeks (for retrograde tracing/optogenetics) or three weeks (for anterograde tracing) before subsequent recording and/or transcardial perfusion.

### Histology and cell counting

Mice were anesthetized via intraperitoneal injection of ketamine (120 mg/kg) and dexmedetomidine (1 mg/kg), then perfused with phosphate-buffered saline (PBS) followed by 4% paraformaldehyde (PFA) in PBS. Brains were extracted and immersed in 4% PFA overnight then in 30% sucrose in PBS for at least 36 hours at 4°C. Coronal brain sections (60 μm thickness) were cut on a cryostat (Leica CM1860) at -25°C and mounted on slides using Invitrogen ProLong Antifade Mountant with DAPI. Fluorescent images were acquired using a slide scanning microscope (Zeiss Axio Scan.Z1) under 20x magnification and ZEN 2.3 software. Whole brain fluorescent images were manually aligned with the reference mouse brain atlas (Franklin & Paxinos) to confirm the accuracy of injection sites and extent of viral spread, and to determine regions of interest (ROIs) for cell counting. For the anterior Pir, four coronal sections between 1.0-2.0 mm anterior to Bregma were counted in each mouse. For the posterior Pir, four coronal sections between 1.5-2.5 mm posterior to bregma were counted in each mouse. Fluorescent-labeled cells within ROIs were manually counted using ImageJ software. For ACx subregions, eight coronal sections between 2.2-3.6 mm posterior to bregma were counted in each mouse. A custom-written MATLAB script was used to register the normalized depth of each cell from the pia during manual counting. Specifically, for each ACx subregion, a line segment spanning the depth of the cortex was drawn perpendicular to the pia in the middle of the subregion. The normalized depth for each cell was defined as the relative length of the projection of the cell’s location onto that line segment.

### Electrophysiological recordings

Recordings were performed in a custom-built acoustic isolation booth. At least one week following headplate surgery, mice were habituated to the recording setup for increasing durations (5 – 60 min) over the course of 3 days. On the day of recording, mice were anesthetized with 1.5-2.5% isoflurane, and a small craniotomy was made over the left auditory cortex using a dental drill, centered ∼1.5mm anterior to the lambdoid suture along the caudal end of the squamosal suture. Following the craniotomy, mice were allowed to recover from anesthesia for at least 30 min. Acute extracellular recordings in ACx were performed using a 32-channel silicon probe (A1x32-Poly2-5mm-50s-177, NeuroNexus). Prior to insertion, the probe was coated in a dye (CM-DiI or SP-DiO; Invitrogen) for post hoc visualization of recording sites. The probe was positioned perpendicularly to the brain surface and lowered at 1-2 μm/s to a final depth of 900-1200 μm or until all channels were within the brain. During lowering of the probe, we monitored responses to brief broadband noise clicks to confirm localization within auditory cortex. Neuronal signals were amplified and digitized with an Intan headstage (RHD 32ch) and recorded by an OpenEphys acquisition board (Siegle et al., 2017) at a rate of 30 kHz. Signals were filtered between 500 and 6000 Hz, offset corrected, and re-referenced to the median across all channels. Spiking data were sorted using Kilosort2 (Pachitariu et al., 2016) and manually curated in phy2 to identify putative single- and multi-units.

### Stimuli

Stimuli during electrophysiological recordings consisted of combinations of auditory and olfactory stimuli. Odors consisted of acetophenone (CAS 98-86-2; Sigma-Aldrich), eucalyptus (100% pure eucalyptus oil; NOW essential oils), or an acetophenone + eucalyptus mixture (1:1 vol/vol), delivered using a respiration-triggered flow dilution olfactometer (Shusterman et al., 2011; Dewan et al., 2018; Williams and Dewan, 2020). We chose these odorants due to their use in previous studies (acetophenone in Cohen et al., 2011), and initially to compare the effects of a monomolecular odorant (acetophenone) versus a more complex mixture (eucalyptus). Because preliminary experiments showed no odor-dependent difference in effects, we combined the groups for subsequent experiments and analysis. This approach allowed us to vary odors between successive recordings in the same mouse, to minimize potential habituation to a single odor. Odorants were liquid diluted 1:1 vol/vol in mineral oil (CAS 8042-47-5; Sigma-Aldrich) in amber vials, then further air diluted (2% N_2_/Air) in the olfactometer, for a final dilution of 1%. The control (non-odorant) vial contained pure mineral oil. The olfactometer and stimulus delivery were controlled by custom Python software and Arduino MEGA 2560. Throughout the recording, we monitored the respiratory cycle of the mouse. Respiration (sniffing) was measured non-invasively using a pressure sensor connected via tubing to the odor port around the mouse’s nose. The pressure was transduced using the sensor and a preamplifier, and the analog signal from the preamplifier was recorded along with electrophysiological data in OpenEphys. Between each trial, the olfactometer tubing was cleared and primed with the next odor for at least 3 s. At the onset of a trial, the mouse’s inhale triggered the ‘final valve’ to switch from delivering continuous clean background air to odorized air, which rapidly delivered the odorized air to the mouse’s nose with no change in total air flow (valve opening duration 1.25 s). Opening of the final valve also triggered playing of the sound stimulus with a delay of 250 ms to allow adequate time for the odor to reach the mouse’s nose and subsequent inhale. The onset timing and kinetics of odors were verified using a photo-ionization detector (miniPID; Aurora Scientific). Sounds consisted of 1 s duration broadband gaussian white noise filtered between 3-60 kHz, either unmodulated or amplitude-modulated (10 Hz, 100% modulation depth). Sounds were generated in Python and converted to analog signals via National Instruments card (NI PCIe-6353, 200 kHz sample rate) and delivered through a speaker (MF-1; Tucker-Davis Technologies) pointed toward the mouse’s right ear, at 0, 45, 55, 65, or 75 dB SPL. Background noise from the olfactometer air flow was measured to be ∼50 dB SPL (1/4-inch condenser microphone; Brüel &Kjær). During recordings, on half of trials, we delivered either odorized air or control air. For recordings with optogenetics, on half of trials, we delivered 470 nm LED light (M470F4; ThorLabs) through a fiber optic cannula (ThorLabs CFML52L05; power 2.8-3.7 mW at cannula tip). LED stimulus onset and duration (1.25 s) were the same as final valve opening. For all recordings, trial order was randomized and different for each recording.

### Electrophysiological data analysis

Neuronal spiking data from each recorded unit were organized by trial type and aligned to the trial onset (i.e. the time of final valve opening). For each unit, peristimulus time histograms (PSTHs) of neuronal spikes were generated using 10 ms time bins, then averaged across trials for each condition. We included all units with baseline firing rate > 1 Hz. For responses to unmodulated noise stimuli, we analyzed spiking during a response window of 0-250 ms after sound onset; for responses to 10 Hz AM stimuli, we analyzed spiking during a response window of 0-1 s. To classify each unit as sound-responsive or odor-modulated, we input the number of spikes during each trial’s response window into a generalized linear model (GLM; predictor variables: sound level [continuous variable normalized from 0-1], odor [0 or 1]; response variable: number of spikes during response window; Poisson distribution, log link function). This allowed the classification of each unit’s responses as having a main effect (*p*<0.05) of sound, odor, and/or a sound-odor interaction. Sound-responsive units that were also responsive to odor or had a significant sound-odor interaction were classified as ‘odor-modulated’ units. For each odor-modulated unit, to determine the direction and magnitude of olfactory modulation, we calculated an Odor Modulation Index (OMI) according to:

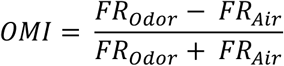

Where FR_Odor_ and FR_Air_ are the mean firing rates in response to sound, with or without odor, respectively. For each unit we calculated the OMI using the mean firing rates for all responses to 5, 15, and 25 dB SNR sounds. Odor-modulated units with OMI>0 were classified as ‘enhanced’ units; units with OMI<0 were classified as ‘suppressed’ units. In addition, for both enhanced and suppressed units, we calculated the OMI for each sound level using the responses to each sound stimulus separately. To calculate z-scored firing rates, we used the mean and standard deviation across trials of firing rate during the 1 s baseline period prior to final valve opening. For ‘enhanced’ units in optogenetics experiments (stGtACR1 and control mice), we calculated an LED Modulation Index according to:

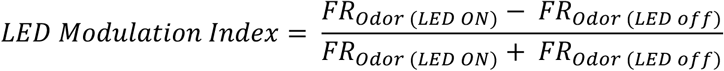

Where FR_Odor (LED ON)_ and FR_Odor (LED off)_ are the mean firing rates in response to Odor+Sound, with or without LED light, respectively.

### Sniff data analysis

Raw analog sniff data from each recording were bandpass filtered between 0.5 and 20 Hz, then subsequent analyses were performed on the filtered sniff traces for each trial during the 1.25 s trial duration, as described above. To calculate sniff frequency for each trial, we measured the average time between peaks (exhales) and between troughs (inhales) in the sniff trace. To calculate sniff amplitude for each trial, we measured the difference between the average peak amplitude minus the average trough amplitude.

### Statistics

Data analysis and figure generation were performed using Matlab or Python. We used the Shapiro-Wilk test to assess normality. For parametric data we used two-tailed paired or unpaired t-tests between groups; for non-parametric data we used two-tailed Wilcoxon signed-rank or Mann-Whitney tests between groups, or Friedman tests for multiple groups. We performed Bonferroni corrections for multiple comparisons. Significance levels are defined as *p*<0.05. Group data in figures are presented as mean ± SEM, unless otherwise noted. The statistical tests performed are indicated in the text and figure legends. Table 1 contains a detailed summary of statistical tests and comparisons.

**Table 1.**
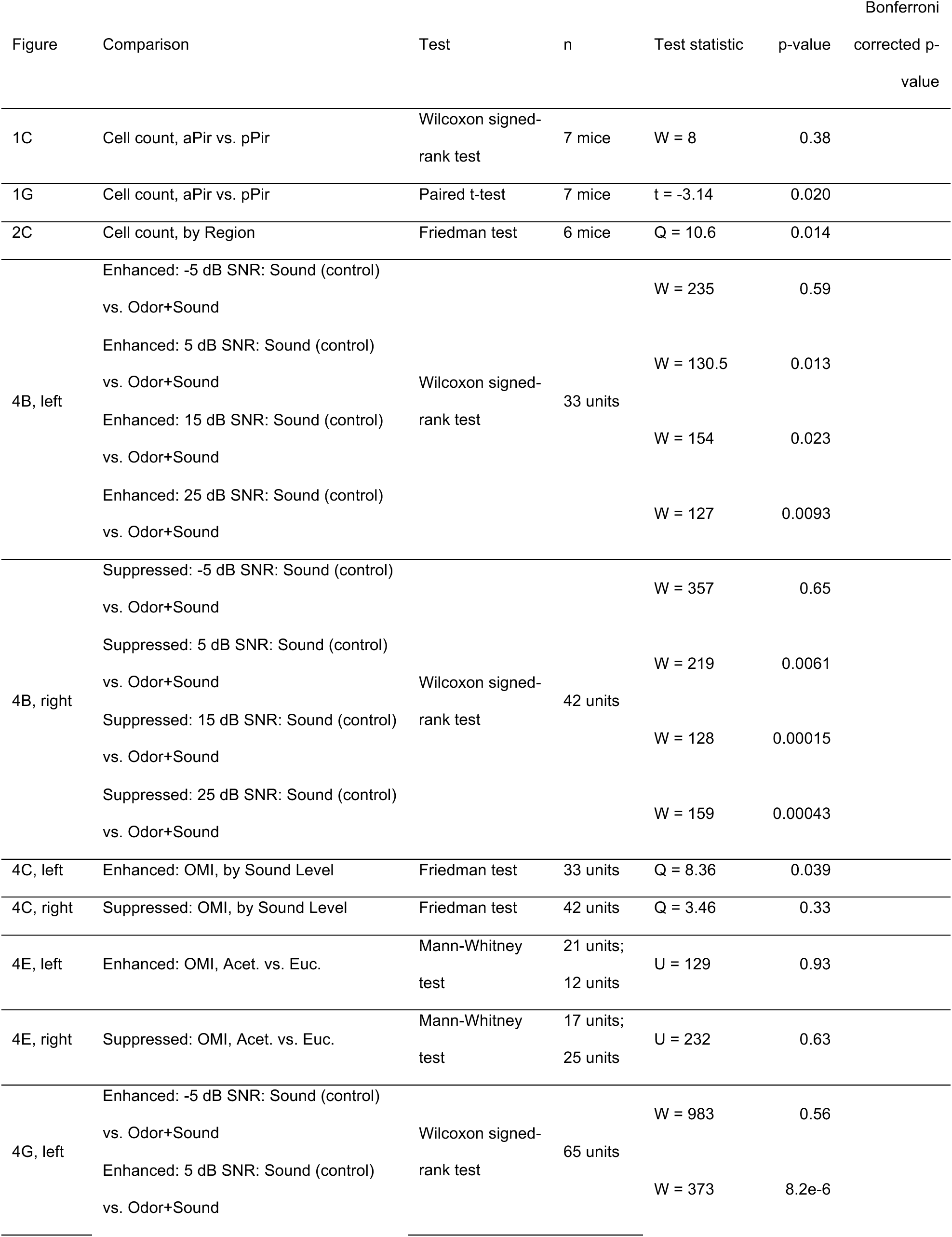
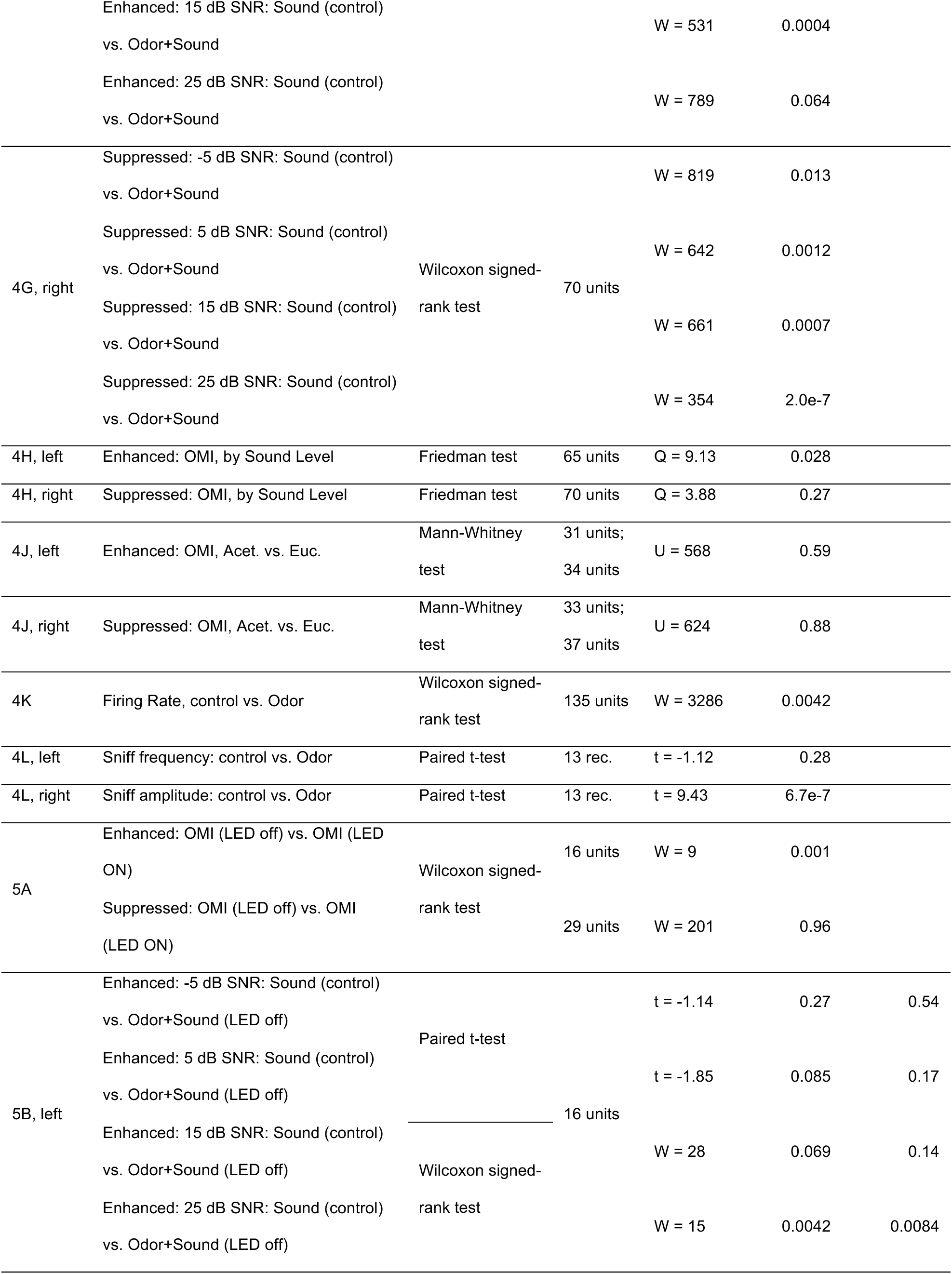
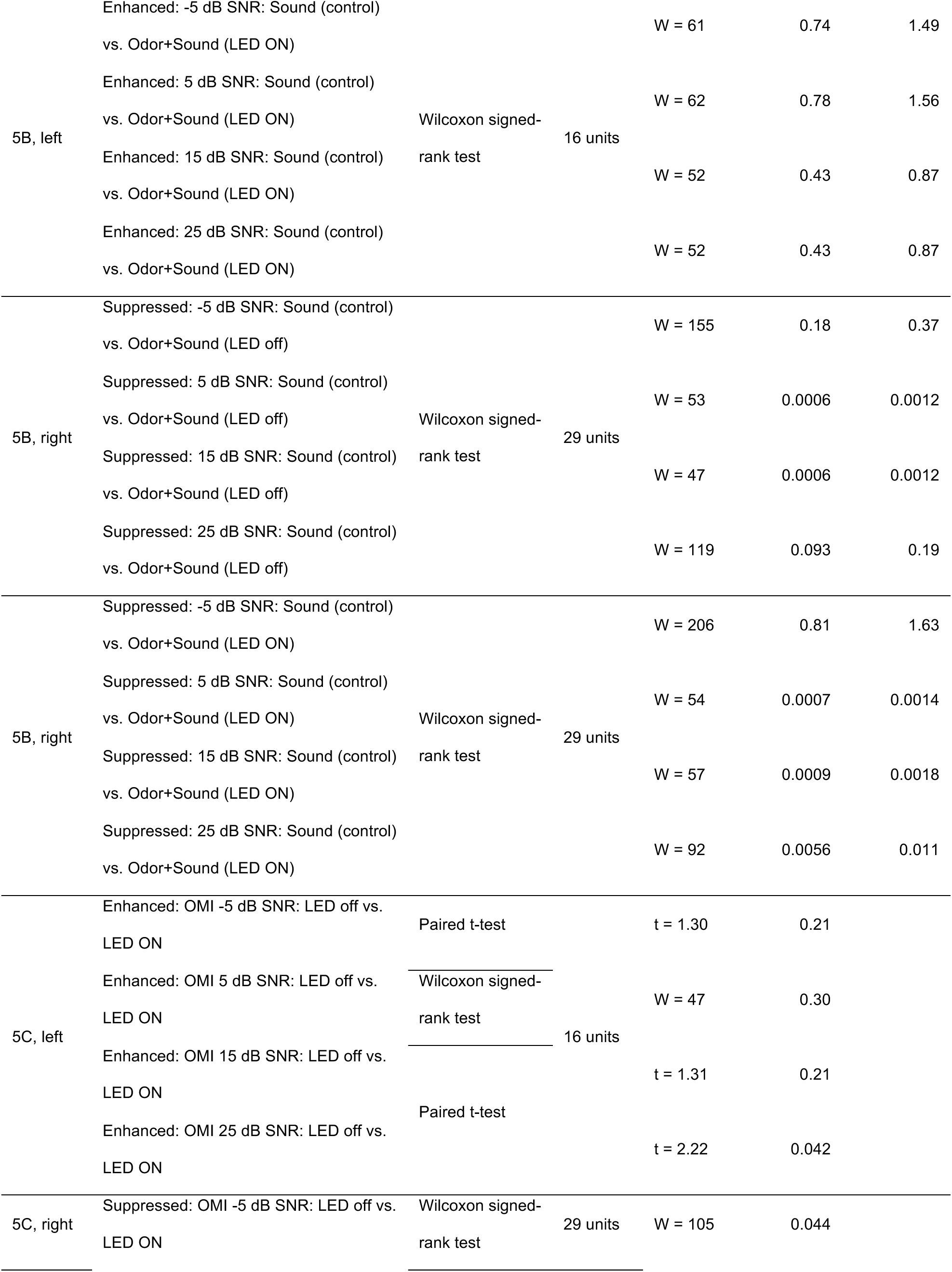
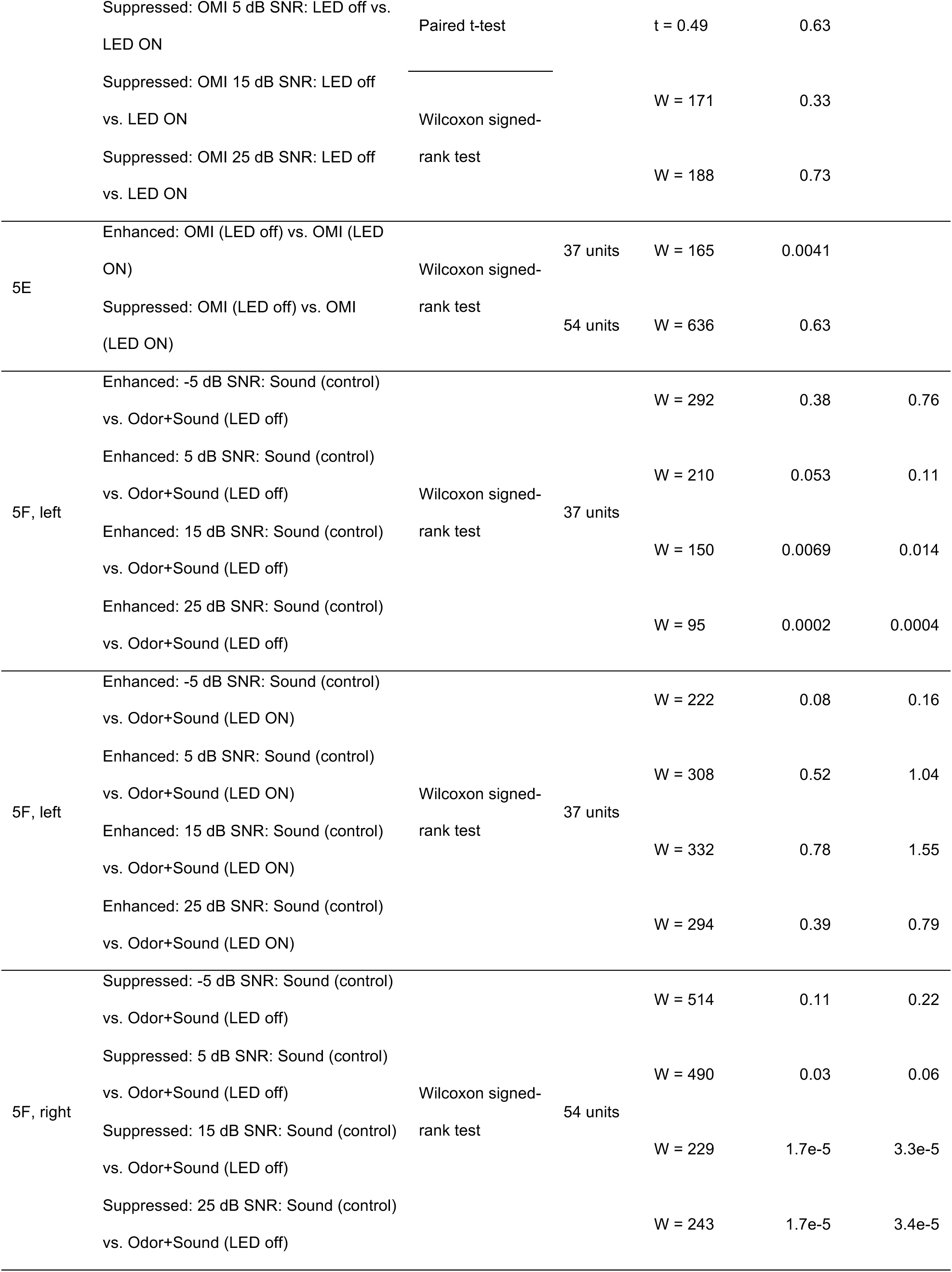
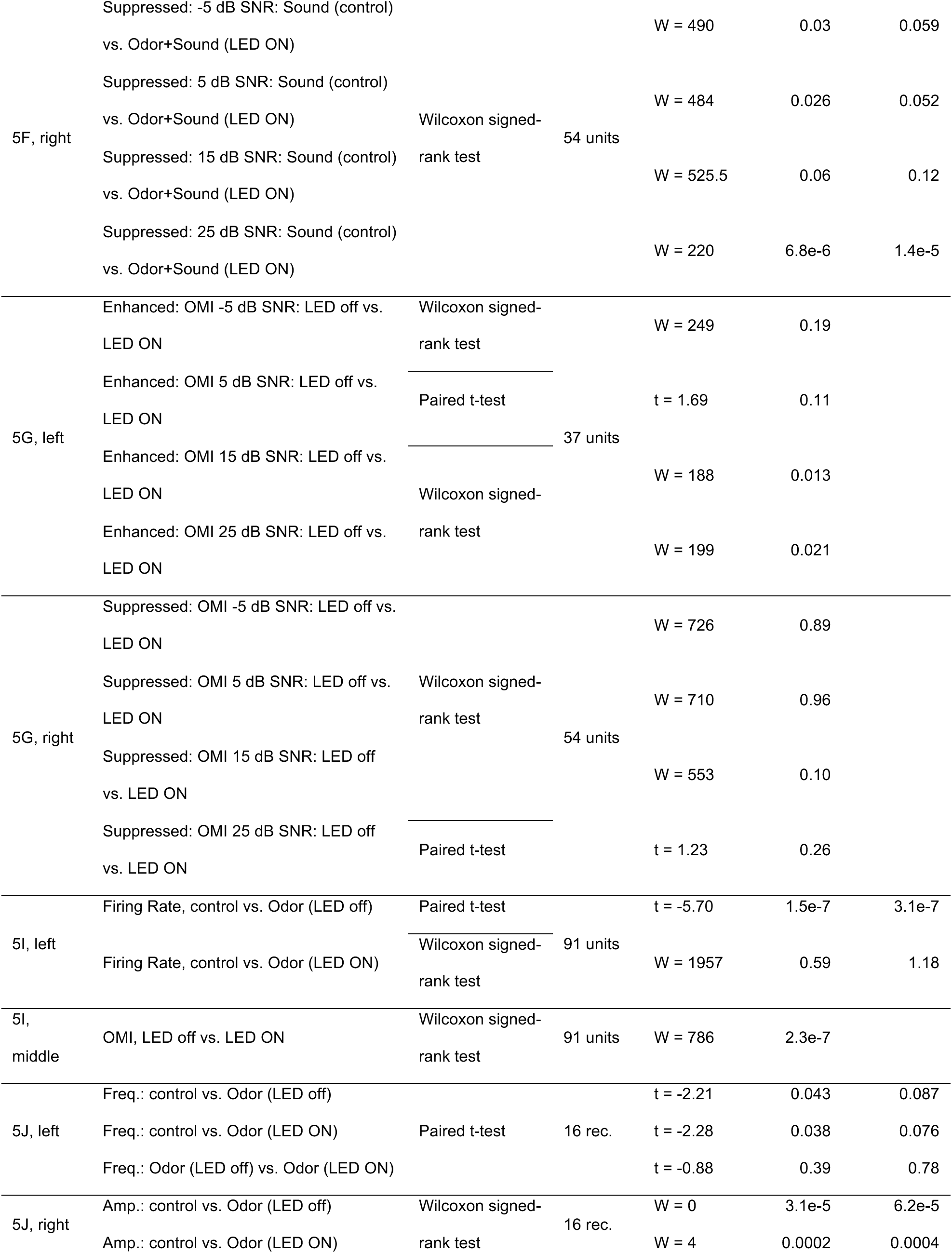
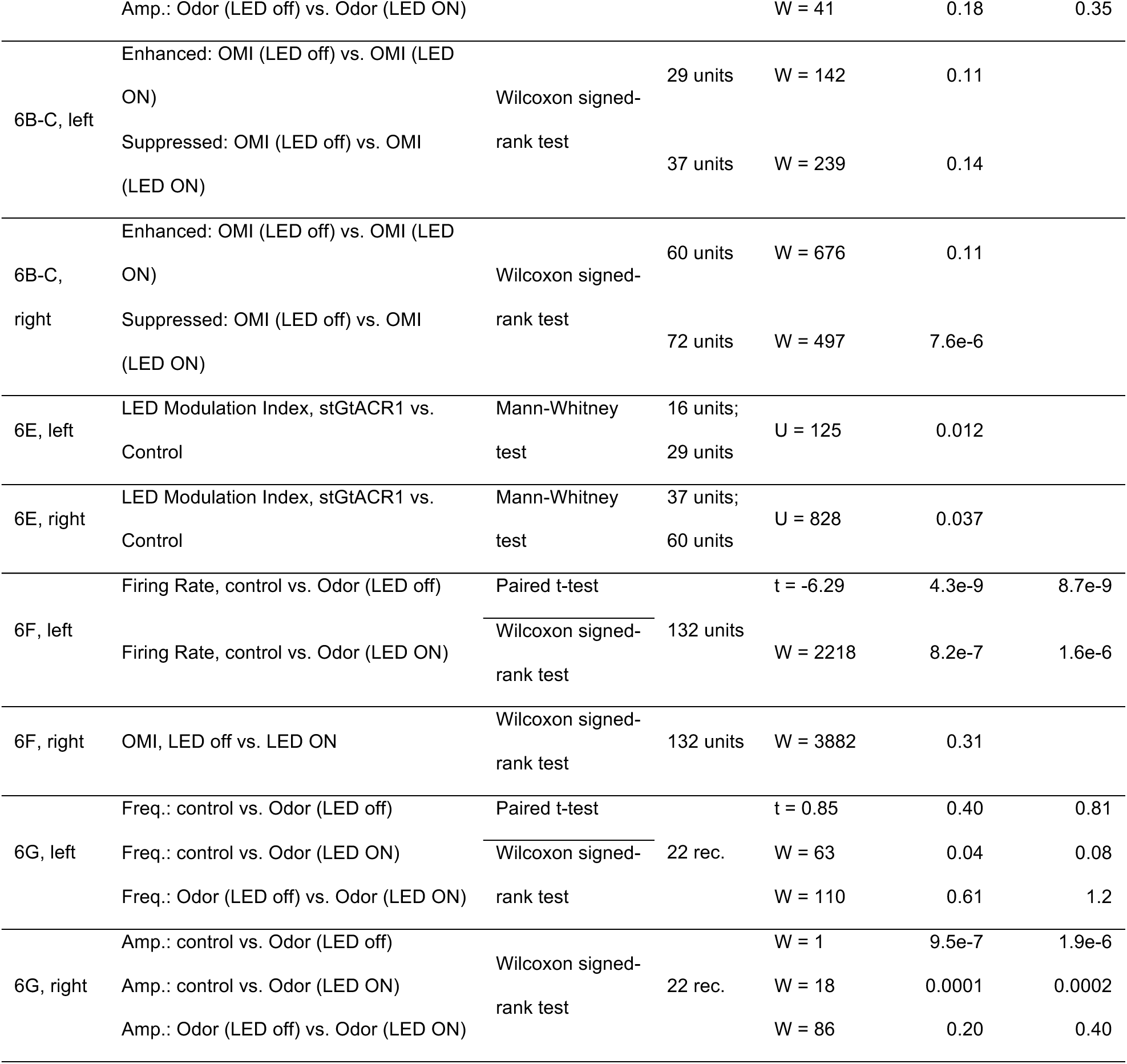
Statistical comparisons. Detailed values and statistical tests for all data figures.

### Data and code availability

Source data is available on Dryad: https://doi.org/10.5061/dryad.dncjsxm7c. Original code is available on GitHub: https://github.com/geffenlab/Vogler_2024.

## Results

### The auditory cortex receives direct input from the piriform cortex

To examine potential anatomical substrates for auditory-olfactory integration in the auditory cortex (ACx), we first sought to identify whether ACx receives direct projections from olfactory brain areas such as the piriform cortex (Pir). We started by injecting a retrograde virus expressing GFP (AAVrg-hSyn-eGFP) into the left ACx of C57BL/6J or Cdh23 mice to selectively label the cell bodies of ACx-projecting neurons (Fig. 1A). We observed fluorescently labeled cells in the ipsilateral Pir (GFP, Fig. 1B). Because the anterior and posterior regions of the Pir are known to exhibit differential long-range connectivity patterns (Bekkers and Suzuki, 2013), we quantified and compared retrograde GFP expression in the anterior Pir (aPir) versus posterior Pir (pPir) (Fig. 1B-C). In most mice, we observed a greater number of labeled cells in pPir compared to aPir. (Fig. 1C), suggesting that the Pir-to-ACx projection predominantly arises from pPir. To confirm the accuracy of injection sites, we examined the extent of viral spread in each mouse (Fig. 1D-E), ensuring localization of the virus injection to ACx with minimal spread into surrounding non-auditory regions.

**Figure 1.**
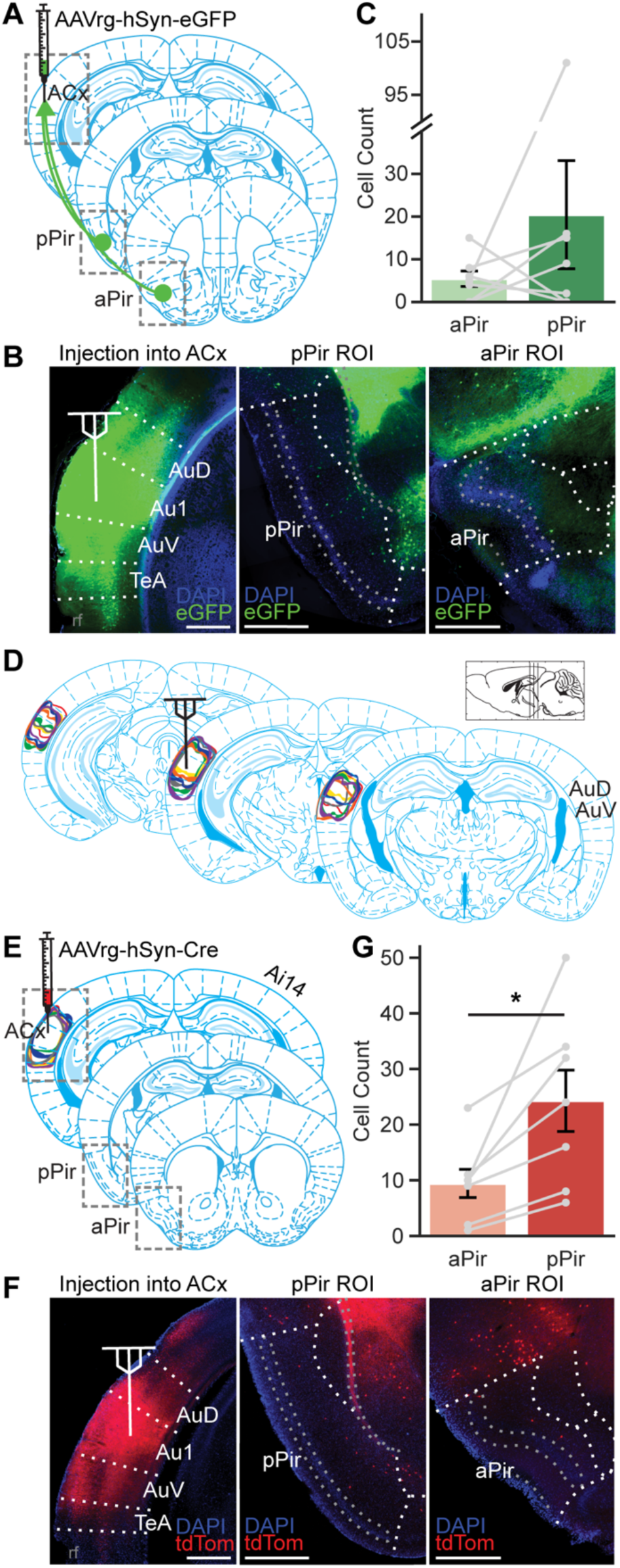
Neurons in the ipsilateral piriform cortex directly project to the auditory cortex. **A)** Injection of AAVrg-hSyn-eGFP into the left auditory cortex (ACx) of C57BL/6J mice (n=5) or Cdh23 mice (n=2). **B)** Representative fluorescent micrographs of ACx injection site (left), posterior piriform (pPir) ROI (middle), anterior piriform (aPir) ROI (right). Scale bars = 500 µm. **C)** Number of cells labeled with GFP in aPir and pPir (n=7 mice; n.s. Wilcoxon signed-rank test). **D)** Viral spread around ACx injection site. Colored outlines indicate extent of viral spread for each mouse (n=7 mice). From left to right, atlas overlays are bregma -3.08 mm, -2.54 mm, and -2.06 mm. **E)** Injection of AAVrg-hSyn-Cre into the left ACx of Ai14 mice (n=7). Colored outlines indicate extent of viral spread around each ACx injection site (n=7 mice). **F)** Representative fluorescent micrographs of ACx injection site (left), pPir ROI (middle), aPir ROI (right). Scale bars = 500 µm. **G)** Number of cells labeled with tdTomato in aPir and pPir (n=7 mice; *p<0.05, paired t-test).

To further confirm the existence of this projection and validate our approach, we performed a similar retrograde tracing experiment in Ai14 cre-reporter mice (progeny of Cdh23 mice crossed with Ai14 mice), injecting retroAAV expressing Cre (AAVrg-hSyn-Cre) into the ACx (Fig. 1E). Again, we found retrogradely labeled cell bodies in ipsilateral Pir (tdTomato, Fig. 1F), and we observed a significantly greater number of labeled cells in pPir compared to aPir (Fig. 1G, *p=0.020, paired t-test). These retrograde tracing experiments reveal a direct projection to ACx arising from Pir, predominantly from pPir.

Next, to further characterize Pir input to ACx, we performed anterograde viral tracing to reveal the targets of Pir neurons within ACx. We injected a transsynaptic anterograde virus (Zingg et al., 2020) expressing Cre (AAV1-hSyn-Cre) into the left Pir of Ai14 cre-reporter mice (progeny of Cdh23 mice crossed with Ai14 mice) to selectively label neurons in ACx that receive Pir input (Fig. 2A). Because our retrograde tracing experiments (Fig. 1) showed that the Pir-to-ACx projection arises mostly from pPir, we specifically targeted pPir for the anterograde virus injections.

**Figure 2.**
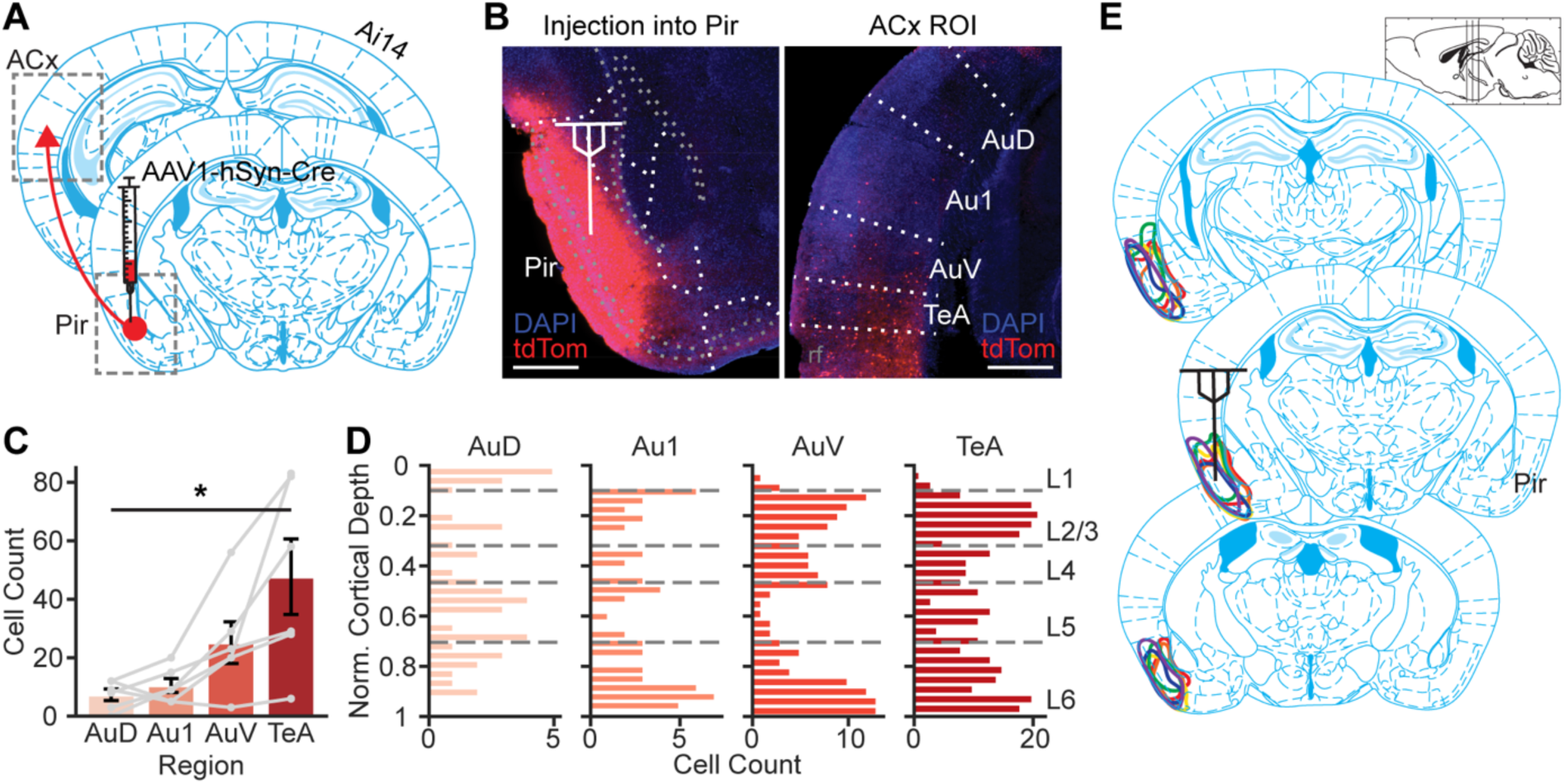
Neurons that receive piriform cortical input span auditory cortical regions. **A)** Injection of AAV1-hSyn-Cre into the left piriform cortex (Pir) of Ai14 mice (n=6). **B)** Representative fluorescent micrographs of the Pir injection site (left) and auditory cortical ROIs (right). Scale bars = 500 µm. **C)** Number of cells labeled with tdTomato across auditory cortical regions (n=6 mice; *p<0.05, Friedman test). **D)** Histograms of the depth distribution (normalized cortical depth) of labeled cells in each auditory cortical region (all cells from 6 mice). **E)** Viral spread around Pir injection site. Colored outlines indicate extent of viral spread for each mouse (n=6 mice). From top to bottom, atlas overlays are bregma -1.94 mm, -1.46 mm, and -0.94 mm.

Consistent with our retrograde tracing experiments, we observed fluorescently labeled cell bodies in the ipsilateral ACx, including the primary ACx (Au1) and the dorsal and ventral secondary ACx (AuD, AuV) (Fig. 2B-C). We additionally observed Pir-recipient cells in the temporal association cortex (TeA) (Fig. 2B-C), consistent with previous studies (Tasaka et al., 2020). We quantified and compared anterograde tdTomato expression across these cortical subregions and found an increase in the number of labeled cells from dorsal-to-ventral auditory subregions (Fig. 2C, *p=0.014, Friedman test). Furthermore, we quantified the cortical depth of the labeled cells and found that Pir-recipient cells span all layers of ACx, but predominantly in layers 2/3 and 6 (Fig. 2D). To again confirm the accuracy of injection sites, we examined the extent of viral spread in each mouse (Fig. 2E), ensuring localization of the virus injection to Pir with minimal spread into surrounding regions. These anterograde tracing experiments further confirm the direct projection from Pir to ACx and reveal that Pir-recipient ACx neurons span auditory cortical and layers and subregions.

Taken together, our tracing experiments confirm a direct projection from pPir to ACx using multiple retrograde and anterograde viral approaches. Our results are consistent with previous studies showing a sparse but detectable Pir-to-ACx projection (Costa et al., 2017; Tasaka et al., 2020). These results suggest an anatomical substrate by which olfactory sensory information may affect auditory processing in ACx, pointing toward a possible mechanism for auditory-olfactory integration.

### *In vivo* electrophysiology for assessing auditory-olfactory integration in auditory cortex

Next, we sought to determine whether and how odors modulate auditory processing in the ACx of awake mice. While previous studies have demonstrated olfactory modulation of sound responses in ACx, most of these studies were performed in anesthetized animals using complex odorants and sound stimuli, or in awake animals but influenced by a behavioral task (Cohen et al., 2011; Cohen and Mizrahi, 2015; Gnaedinger et al., 2019; Gilday and Mizrahi, 2023). As such, it remains unclear how auditory-olfactory integration may depend on brain state, stimulus complexity, and/or experience-dependent learning. We therefore sought to answer the basic question of whether and how a temporally coupled odor modulates ACx responses to simple sounds in untrained, awake mice. To answer this question, we designed a custom-made system combining sound stimuli with a respiration-triggered olfactometer to deliver sounds and odors to awake head-fixed mice with high temporal precision during *in vivo* electrophysiology (Fig. 3A). We recorded the activity of ACx neurons in response to combinations of odor (acetophenone or eucalyptus, 1%) and sound stimuli (unmodulated or amplitude-modulated [AM, 10 Hz] noise, 0 – 75 dB). We used the respiration-triggered olfactometer to control the timing of odor/sound stimulus delivery with the mouse’s inhale (Fig. 3B). To examine auditory-olfactory integration in ACx, we targeted recordings to the ACx (Fig. 3C) and recorded the activity of single- and multi-units in response to sounds, with or without odor (Fig. 3D).

**Figure 3.**
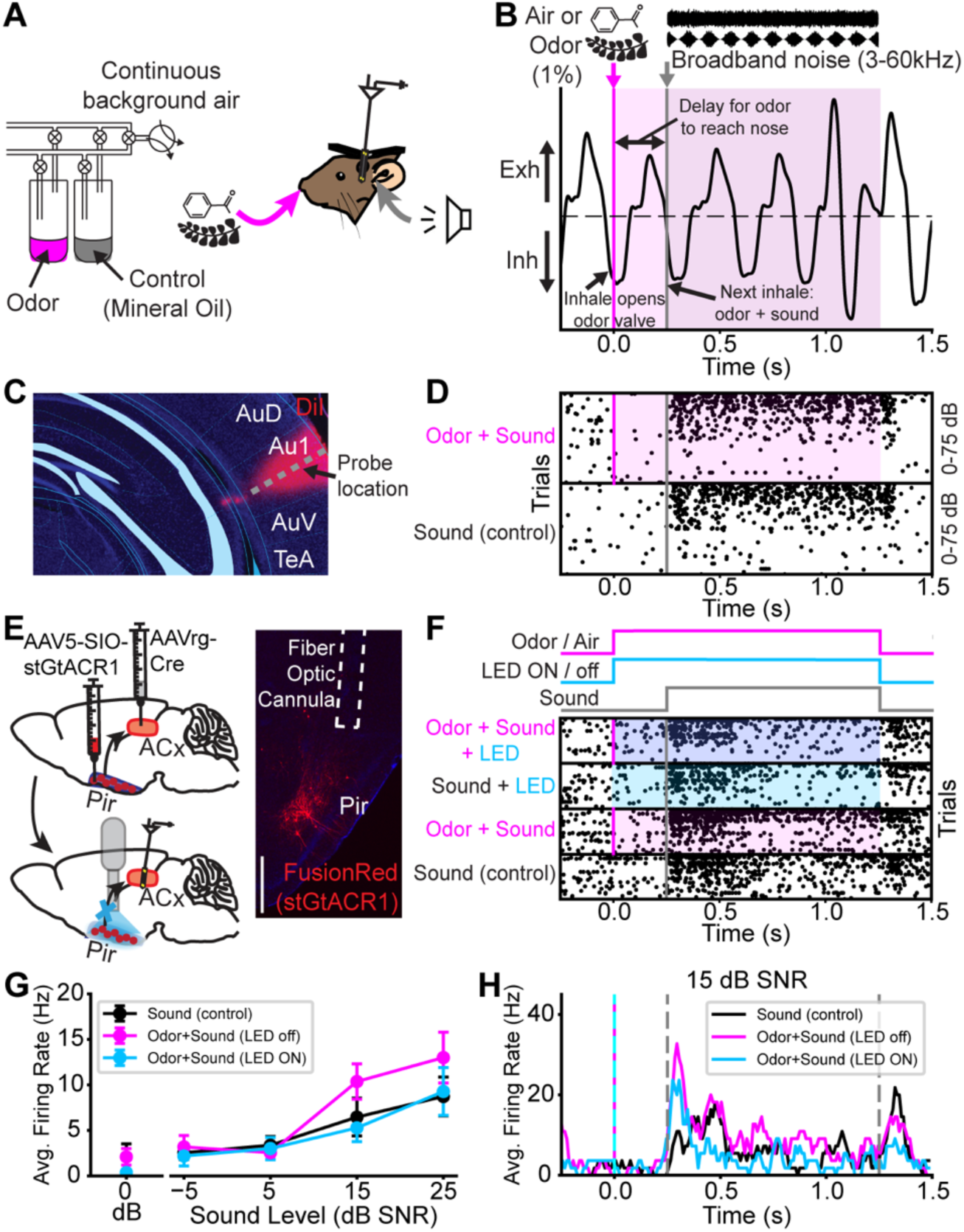
*In vivo* electrophysiology for assessing auditory-olfactory integration in auditory cortex. **A)** Experimental setup for delivering combinations of odor/sounds to awake mice during recordings. **B)** Stimulus timing using respiration-triggered olfactometer. **C)** Example post-hoc visualization of probe location using DiI. **D)** Example raster plot from sound-responsive ACx unit. Trials sorted by sound intensity and odor/control**. E)** Virus injections for selectively expressing stGtACR1 in ACx-projecting Pir neurons in C57BL/6J or Cdh23 mice. Right: example opsin expression in Pir and cannula placement for in vivo optogenetics using 470 nm LED. Scale bar = 500 µm. **F)** Stimulus timing during optogenetic experiments and example raster plot from ACx unit. Trials sorted by sound intensity and odor/LED combination. **G)** Average firing rates (Hz) of example unit shown in F, in response to various sound levels (left: 0 dB [no sound]; right: dB SNR [relative to background noise of olfactometer air flow]) in control and odor, with and without LED inactivation of PCx-to-ACx projections. **H)** Peristimulus time histogram (PSTH) of example unit shown in F and G, in response to 15 dB SNR sound.

Because our anatomical experiments pointed toward the Pir-to-ACx pathway as a possible substrate for auditory-olfactory integration in ACx, we tested whether and how this pathway contributes to odor modulation of sound responses using an *in vivo* optogenetic approach. To selectively inactivate ACx-projecting Pir neurons in C57BL/6J or Cdh23 mice, we injected AAVrg-hSyn-Cre into ACx, and the cre-dependent soma-targeted inhibitory opsin stGtACR1 (AAV5-SIO-stGtACR1) (Mahn et al., 2018) into pPir (Fig. 3E). During *in vivo* recordings, we delivered 470 nm LED light through a fiber optic cannula implanted over pPir (Fig. 3E-F). Fig. 3G-H shows sound responses in an example ACx unit under the various trial conditions: odor enhances the sound responses, but this enhancement is eliminated by LED inactivation of ACx-projecting Pir neurons.

### Odor modulates sound responses in auditory cortex

First, we tested whether and how odors modulate responses to sound stimuli in ACx. We identified ‘sound-responsive’ and ‘odor-modulated’ units using a generalized linear model (GLM). In response to unmodulated broadband noise, ∼20% of sound-responsive units were significantly modulated by odor (Fig. 4A). However, the effects of the odor on sound responses were heterogenous: in response to unmodulated noise, we observed both enhancement (Fig. 4B-D, left) and suppression (Fig. 4B-D, right) of sound responses by odor. To quantify the amount of enhancement or suppression in odor-modulated units, we calculated an Odor Modulation Index (OMI) for each unit for each sound level (Fig. 4C). We found that OMIs vary significantly across sound level for enhanced units (Fig. 4C, left; *p=0.039, Friedman test), but not suppressed units (Fig. 4C, right; n.s. p=0.326, Friedman test). Because we used acetophenone and eucalyptus as odorants in our experiments and varied the odor between recordings, we tested for an odor-dependent difference in enhancement or suppression. We compared the amount of odor modulation (OMI) using acetophenone versus eucalyptus (Fig. 4E) and found that odor modulation does not differ between odorants (Fig. 4E, left: n.s. p=0.93, Mann-Whitney test; Fig 4E, right: n.s. p=0.63, Mann-Whitney test). Therefore, we combined the groups here and in subsequent experiments and analyses.

**Figure 4.**
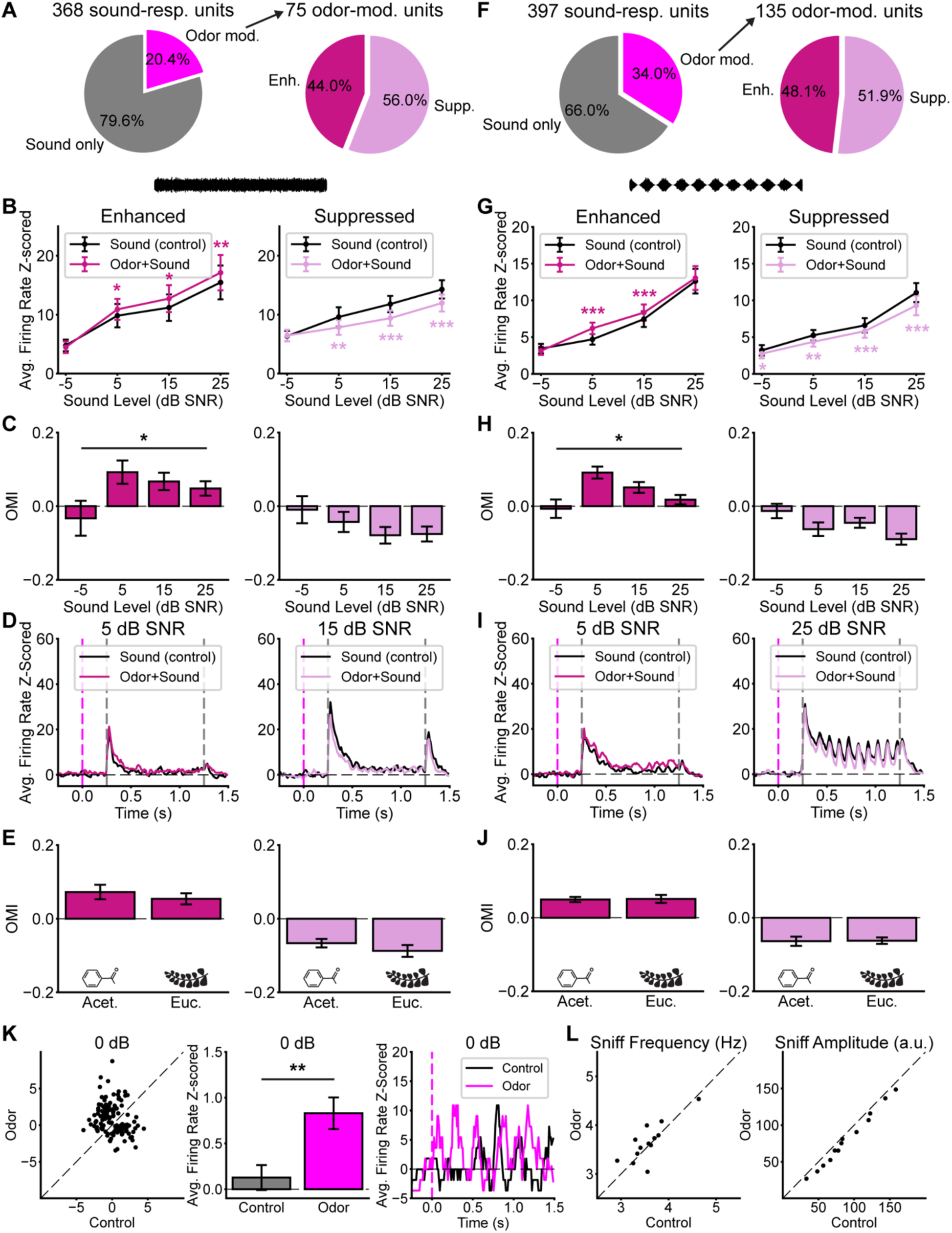
Odor modulates sound responses in auditory cortex. **A)** Left: Percentage of significantly odor-modulated units in ACx in response to unmodulated broadband noise, according to GLM. Right: percentage of Enhanced vs Suppressed odor-modulated units. n=5 mice (1 C57BL/6J, 4 Cdh23), 368 sound-responsive units, 293 ‘Sound only’ units, 75 ‘Odor mod.’ units, 33 Enhanced units, 42 Suppressed units. **B)** Average responses to sound levels (dB SNR relative to noise of background air flow), with Odor versus control, for Enhanced units (left) and Suppressed units (right). *p<0.05, **p<0.01, ***p<0.001, Wilcoxon signed-rank tests. **C)** Average Odor Modulation Index (OMI) for each sound level, for Enhanced units (left) and Suppressed units (right). *p<0.05, Friedman test. **D)** Average PSTHs in response to 5 dB SNR (left) and 15 dB SNR (right) sound. Dashed magenta line indicates odor valve opening time, gray lines indicate sound stimulus start and stop times. **E)** Average OMI for experiments with acetophenone (Acet.) versus eucalyptus (Euc.), for enhanced units (left; Acet. n=21 units, Euc. n=12 units) and suppressed units (right; Acet. n=17 units, Euc. n=25 units). n.s. Mann-Whitney tests. **F-J)** Same as A-E, for 10 Hz amplitude modulated noise. n= 5 mice, 397 sound-responsive units, 262 ‘Sound only’ units, 135 ‘Odor mod.’ units, 65 Enhanced units, 70 Suppressed units. **G)** *p<0.05, **p<0.01, ***p<0.001, Wilcoxon signed-rank tests. **H)** *p<0.05, Friedman test. **J)** Left: Acet. n=31 units, Euc. n=34 units. Right: Acet. n=33 units, Euc. n=37 units. n.s. Mann-Whitney tests. **K)** Left and middle: Average firing rate (z-scored) for odor-modulated units (10 Hz AM; units in F, n=135 units), in response to Odor versus control, in the absence of sound (0 dB). **p<0.01, Wilcoxon signed-rank test. Right: PSTH from an example odor-modulated unit, in the absence of sound (0 dB). **L)** Left: Average sniff frequency during Odor versus control trials (n=13 recordings). n.s. paired t-test. Right: Average sniff amplitude during Odor versus control trials (n=13 recordings). ***p<0.001, paired t-test.

In addition, we tested the effect of odor on responses to 10 Hz amplitude-modulated (AM) noise and performed the same analyses (Fig. 4F-J). In response to AM noise, we found a greater proportion of odor-modulated sound-responsive units (∼33%, Fig. 4F). We again observed both enhancement and suppression of sound responses by odor (Fig. 4G-I), and that OMIs vary significantly across sound level for enhanced units (Fig. 4H, left; *p=0.028, Friedman test). Moreover, we again found that odor modulation does not differ between odorants (Fig. 4J, left: n.s. p=0.59, Mann-Whitney test; Fig 4J, right: n.s. p=0.88, Mann-Whitney test).

In sound-responsive odor-modulated units, we tested the effect of odor on neuronal firing in the absence of sound. We found an overall slight but significant enhancement of neuronal firing rate in response to odor compared to control (Fig. 4K, **p=0.0042, Wilcoxon signed-rank test). This result suggests that odors may impact ACx processing in the absence of sound, as well as in response to sound stimuli. Specifically, odors enhance baseline neuronal firing, but either enhance or suppress sound-evoked responses.

In response to odors, rodents actively change their respiratory patterns (sniffing) (Wesson et al., 2009; Wachowiak, 2011). To evaluate the potential effects of respiration on the observed changes in neuronal firing in our experiments, we compared the sniff frequency and amplitude during odor versus control trials in each recording (Fig. 4L). We observed that while odor did not significantly affect sniff frequency (Fig. 4L, left; n.s. p=0.28, paired t-test), odor reduced sniff amplitude (Fig. 4L, right; ***p=6.7e-7). This may reflect shallower breathing in response to the odor in our experiments. Nevertheless, our results demonstrate odor-evoked changes in ACx activity, enabling further dissection of the underlying mechanisms.

Together, these results demonstrate that: 1) olfactory stimuli modulate auditory processing in awake mice, including enhancement of baseline firing as well as bidirectional modulation of sound-evoked responses; and 2) olfactory stimuli enhance or suppress neuronal responses to multiple types of sound stimuli, including both unmodulated and AM noise.

### Piriform input to auditory cortex contributes to auditory-olfactory integration

To test the role of the newly characterized Pir-to-ACx pathway (Fig. 1-2) in auditory-olfactory olfactory integration, we combined *in vivo* electrophysiology in awake mice with optogenetics specifically targeting ACx-projecting Pir neurons (Fig. 3). We again recorded sound responses with or without odor, but now we also optogenetically inactivated ACx-projecting Pir neurons by shining LED light through a cannula implanted over ipsilateral pPir (Fig. 3E-H). Consistent with our previous experiments (Fig. 4), we observed both enhancement and suppression of sound responses by odor in a subpopulation of ACx units (Fig. 5A). For both the enhanced and suppressed units, we compared the amount of odor modulation, using the OMI, with or without LED-inactivation of Pir neurons. Strikingly, we observed that for the enhanced units, the LED significantly reduced odor modulation (**p=0.001, Wilcoxon signed-rank test), whereas odor modulation in suppressed units was unaffected (n.s. p=0.96, Wilcoxon signed-rank test) (Fig. 5A). We then examined the effect of the LED on odor modulation for each sound level (Fig. 5B-C) and found that in enhanced units, the LED significantly reduced the amount of odor modulation (OMI) at the highest sound level (Fig. 5C, left; 25 dB SNR: *p=0.042, Wilcoxon signed-rank test), whereas in suppressed units the odor modulation was unaffected (Fig. 5C, right; 25 dB SNR: n.s. p=0.73, Wilcoxon signed-rank test). We again repeated the same analyses for responses to AM noise and observed similar results: the LED significantly reduced the amount of odor modulation in enhanced units (**p=0.004, Wilcoxon signed-rank test), but not in suppressed units (n.s. p=0.63, Wilcoxon signed-rank test) (Fig. 5E). For enhanced units, the LED significantly reduced the amount of odor modulation at 15 dB SNR and 25 dB SNR (Fig. 5G, left; 15 dB SNR: *p=0.013; 25 dB SNR: *p=0.021; Wilcoxon signed-rank tests), whereas suppressed units were unaffected (Fig. 5G, right; 15 dB SNR: n.s. p=0.10, Wilcoxon signed-rank test; 25 dB SNR: n.s. p=0.23, paired t-test). These results demonstrate that odor *enhancement* of sound responses, but not odor suppression, requires the Pir-to-ACx pathway, suggesting that this pathway provides an excitatory drive.

**Figure 5.**
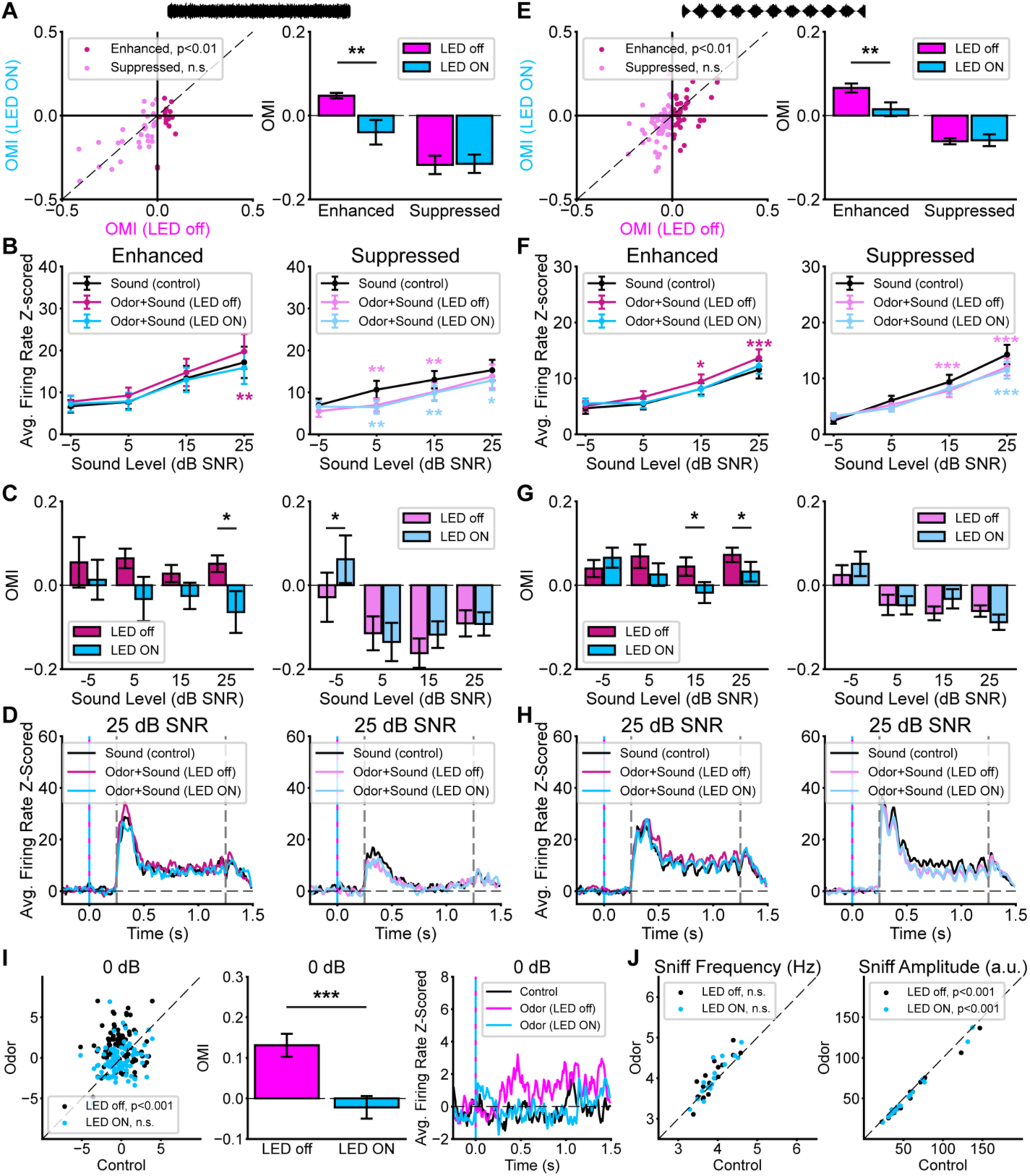
*In vivo* optogenetic inactivation of ACx-projecting piriform neurons reduces odor-mediated enhancement but not suppression. A) Left: Odor Modulation Index (OMI) with vs without LED, for Enhanced and Suppressed ACx units in response to unmodulated broadband noise. n= 6 mice (3 C57BL/6J, 3 Cdh23), 16 Enhanced units, 29 Suppressed units. Wilcoxon signed-rank tests LED off vs LED ON. Right: Average OMI for Enhanced and Suppressed units, LED off vs LED ON. **p<0.01, Wilcoxon signed-rank test. **B)** Average responses to various sound levels (dB SNR relative to noise of background air flow): sound alone versus odor with or without LED, for Enhanced units (left) and Suppressed units (right). *p<0.05, **p<0.01, Wilcoxon signed-rank tests vs. ‘Sound (control)’. **C)** Average OMI for each sound level, for Enhanced units (left) and Suppressed units (right). *p<0.05, Wilcoxon signed-rank test. **D)** Average PSTHs in response to 25 dB SNR sound stimulus. Dashed magenta and cyan lines indicate odor valve opening and LED ON time, gray lines indicate sound stimulus start and stop times. **E-H)** Same as A-D, for 10 Hz amplitude modulated noise. n= 6 mice, 37 Enhanced units, 54 Suppressed units. E) **p<0.01, Wilcoxon signed-rank test. **F)** *p<0.05, ***p<0.001, Wilcoxon signed-rank tests vs. ‘Sound (control)’ **G)** *p<0.05, Wilcoxon signed-rank tests. **I)** Left: Average firing rate (z-scored) for odor modulated units (10 Hz AM; units in E, n=91 units) in the absence of sound (0 dB), in response to Odor versus control with LED off and LED ON. LED off: ***p<0.001, paired t-test. LED ON: n.s. p>1, Wilcoxon signed-rank test. Middle: Average OMI in the absence of sound (0 dB), LED off vs LED ON, ***p<0.001, Wilcoxon signed-rank test. Right: Average PSTH for odor-modulated units, in the absence of sound (0 dB). **J)** Left: Average sniff frequency during Odor versus control trials, with LED off and LED ON (n=16 recordings). LED off: n.s. LED ON: n.s. paired t-tests. LED off vs LED ON: n.s. paired t-test. Right: Average sniff amplitude during Odor versus control trials (n=13 recordings). LED off: ***p<0.001, LED ON: ***p<0.001, Wilcoxon signed-rank tests. Odor (LED off) vs Odor (LED ON): n.s. Wilcoxon signed-rank test.

In addition, we examined odor-driven enhancement of ACx neuronal firing in the absence of sound, with or without *in vivo* optogenetic inactivation of Pir neurons. Consistent with our previous experiments (Fig. 4K), in odor-modulated ACx units we observed that odor significantly enhanced neuronal firing rate, compared to control (Fig. 5I, left; LED off: ***p=3.07e-7, paired t-test). Strikingly, however, this enhancement was eliminated by the LED (Fig. 5I, left; LED ON, n.s. p>1, Wilcoxon signed-rank test). When comparing the amount of odor modulation (OMI) with versus without the LED, we found that the LED significantly reduced the odor-driven enhancement in the absence of sound (Fig. 5I, middle; ***p=2.3e-7, Wilcoxon signed-rank test). This result suggests that optogenetic inactivation of ACx-projecting Pir neurons eliminates odor-driven enhancement of baseline ACx neuronal firing, and therefore that this enhancement requires the direct Pir-to-ACx pathway.

Finally, we examined potential effects of respiration on the changes in neuronal firing we observed in our optogenetic experiments (Fig. 5J). Consistent with our previous results (Fig. 4L), we observed an odor-mediated reduction of sniff amplitude (Fig. 5J, right; LED off, ***p=6.2e-5, Wilcoxon signed-rank test) but not sniff frequency (Fig. 5J, left; LED off, n.s. p=0.087, paired t-test). Importantly, during trials with the LED, we observed similar effects (Fig. 5J, right: LED ON, ***p=0.0004, Wilcoxon signed-rank test; Fig. 5J, left: LED ON, n.s. p=0.076, paired t-test). Moreover, when comparing respiration during odor trials with versus without the LED, we observed that the LED did not affect sniff frequency (Fig. 5J, left; n.s. p=0.78, paired t-test) or amplitude (Fig. 5J, right, n.s. p=0.35, Wilcoxon signed-rank test). These results critically demonstrate that the optogenetic reduction of odor-driven enhancement is not due to any effects of the LED on sniffing, and is likely specifically due to inactivation of the Pir-to-ACx pathway.

To further confirm that the optogenetic effects we observed (Fig. 5) were specifically due to inactivation of Pir-to-ACx input via light activation of stGtACR1, and not potential other non-specific effects of the LED light, we performed additional control experiments. Control Cdh23 mice underwent an identical experimental procedure as the optogenetic mice, except we injected a control virus into Pir (AAV5-Flex-tdTomato) instead of opsin (Fig. 6A). In control mice, LED light did not significantly reduce odor modulation in enhanced units, for both unmodulated noise (Fig. 6B-C, left; n.s. p=0.11, Wilcoxon signed-rank test) and AM noise (Fig. 6B-C, right; n.s. 0.115, Wilcoxon signed-rank test). Next, we examined the effect of the LED on enhanced units in the controls. To quantify the effect of the LED on olfactory modulation, we calculated an LED Modulation Index (Fig. 6E). For both unmodulated noise (Fig. 6E, left; **p=0.01, Mann-Whitney test) and AM noise (Fig. 6E, right; *p=0.037, Mann-Whitney test), the LED significantly reduced odor modulation in mice expressing stGtACR1 as compared to controls.

**Figure 6.**
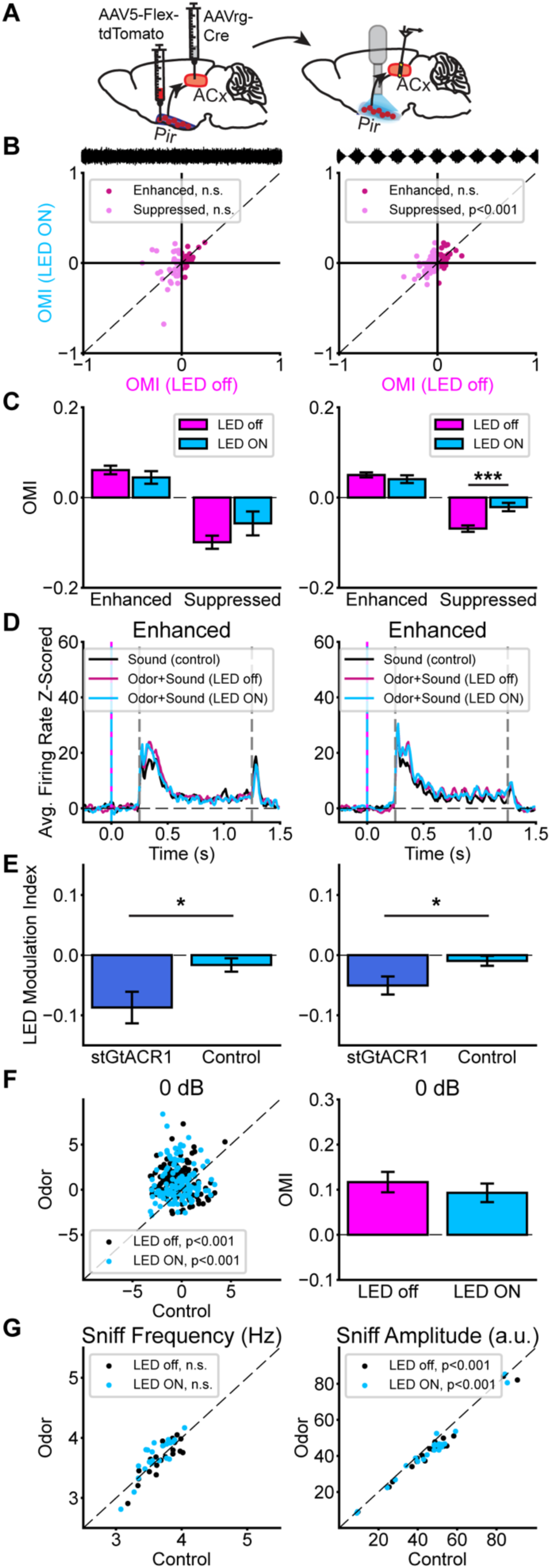
Optogenetic inhibition of odor-mediated enhancement is specifically due to inactivation of ACx-projecting Pir neurons. **A)** Virus injections for optogenetic control experiments expressing control virus in ACx-projecting Pir neurons in Cdh23 mice. **B)** Left: OMI with vs without LED, for Enhanced and Suppressed ACx units in response to unmodulated broadband noise. n= 7 Cdh23 mice, 29 Enhanced units, 37 Suppressed units. Wilcoxon signed-rank tests LED off vs LED ON. Right: Same as Left, but in response to 10 Hz AM noise. n= 7 mice, 60 Enhanced units, 72 Suppressed units. Wilcoxon signed-rank tests LED off vs LED ON. **C)** Average OMI for Enhanced and Suppressed units, LED off vs LED ON., in response to unmodulated noise (left) and 10 Hz AM noise (right). ***p<0.001, Wilcoxon signed-rank test. **D)** Average PSTHs in response to 25 dB SNR sound, for unmodulated (left) and 10 Hz AM noise (right). **E)** LED Modulation Index for Enhanced units in response to unmodulated (left) and 10 Hz AM (right) noise, in stGtACR1 mice versus Control mice. Left: stGtACR1: n=16 units; Control: n=29 units; *p<0.05, Mann-Whitney test. Right: stGtACR1: n=37 units; Control: n=60 units; *p<0.05, Mann-Whitney test. **F)** Left: Average firing rate (z-scored) for odor-modulated units (10 Hz AM; units in B, right, n=132 units) in the absence of sound (0 dB), in response to Odor versus control with LED off and LED ON. LED off: ***p<0.001, paired t-test. LED ON: ***p<0.001, Wilcoxon signed-rank test. Right: Average OMI in the absence of sound (0 dB), LED off vs LED ON. n.s. Wilcoxon signed-rank test. **G)** Left: Average sniff frequency during Odor versus control trials, with LED off and LED ON (n=22 recordings). LED off: n.s. paired t-test, LED ON: n.s. Wilcoxon signed-rank test. LED off vs LED ON: n.s. Wilcoxon signed-rank test. Right: Average sniff amplitude during Odor versus control trials (n=22 recordings). LED off: ***p<0.001, LED ON: ***p<0.001, Wilcoxon signed-rank tests. Odor (LED off) vs Odor (LED ON): n.s. Wilcoxon signed-rank test.

In control mice we again observed odor-driven enhancement of baseline neuronal firing in the absence of sound (Fig. 6F, left; LED off, ***p=8.7e-9, paired t-test); however, this enhancement persisted with the LED (Fig. 6F, left; LED ON, ***p=1.64e-6, Wilcoxon signed-rank test). When comparing the amount of odor modulation (OMI) with versus without the LED, we found that the LED did not affect the odor-driven enhancement in the absence of sound (Fig. 6F, right; n.s. p=0.31, Wilcoxon signed-rank test). Additionally, when comparing respiration during the various stimulation conditions, we again confirmed that the LED did not affect sniff frequency (Fig. 6G, left; n.s. p>1, Wilcoxon signed-rank test) or amplitude (Fig. 6G, right; n.s. p=0.40, Wilcoxon signed-rank test). These results confirm that the reduction of odor-driven enhancement we observed in mice expressing stGtACR1 (Fig. 5) was specifically due to inactivation of the Pir-to-ACx pathway.

Together, our results identify the role of the Pir-to-ACx pathway in auditory-olfactory integration. Optogenetic inactivation of ACx-projecting Pir neurons selectively reduced odor-driven enhancement of baseline ACx firing as well as enhancement of sound-evoked responses. Therefore, the Pir-to-ACx pathway contributes to odor-mediated enhancement of sound responses, but not suppression.

## Discussion

In this study we tested the functional role of a direct projection from the piriform cortex to the auditory cortex. We identified the Pir-to-ACx pathway using retrograde and anterograde viral tracing, highlighting that this pathway arises primarily from posterior Pir (Fig. 1), and projects to multiple ACx subregions and laminae (Fig. 2). Next, using *in vivo* electrophysiology in ACx, we demonstrated that olfactory stimuli enhance baseline activity in ACx neurons and bidirectionally modulate sound-evoked activity, either enhancing or suppressing sound responses (Figs. 3-4). Finally, using *in vivo* optogenetics during electrophysiology, we found that the Pir-to-ACx projection is necessary for odor-mediated enhancement of ACx activity and sound responses, but is minimally engaged during suppression (Figs. 5-6). Our results reveal that the Pir-to-ACx pathway shapes auditory-olfactory integration in ACx, contributing to odor-mediated enhancement of sound processing.

Our anatomical results demonstrate that the Pir-to-ACx projection primarily arises from posterior Pir (pPir), and projects across multiple ACx subregions and cortical layers. Its origination in pPir is consistent with previous studies showing that pPir contains more associational connections with other higher-order brain areas compared to anterior Pir (Bekkers and Suzuki, 2013). For example, pPir is known to have interconnections with the prefrontal cortex, amygdala, and entorhinal and perirhinal cortices, and is thought to encode odor quality (Johnson et al., 2000; Zelano et al., 2011). Our results add ACx to the list of Pir-recipient regions and expand knowledge of cross-modal cortical circuitry. Moreover, the distribution of Pir-recipient neurons spanning ACx is consistent with previous studies showing that cortico-cortical intra-telencephalic neurons project to/from multiple cortical layers including L2/3 and L6 (Shepherd, 2013; Harris and Shepherd, 2015).

Notably, while previous studies have shown evidence for the Pir-to-ACx projection (Costa et al., 2017; Tasaka et al., 2020), such findings have gone relatively underexplored. Our results are consistent with these previous studies and provide further evidence for the existence and functional impact of the Pir-to-ACx cross-cortical circuit. Although this projection is sparse, we revealed this circuity using a rigorous approach comprised of four different viral strategies: retroAAV-GFP (Fig. 1), retroAAV-Cre (Fig. 1), AAV1-Cre (Fig. 2), and selective labeling of ACx-projecting Pir neurons for optogenetics (retroAAV-Cre and AAV5-stGtACR1, Fig. 3). It is possible that a few Pir cells labeled as ACx-projecting (Fig. 1) or Pir-recipient ACx cells (Fig. 2) may instead result from minor viral spread outside the target region (Fig. 1D-E, 2E); however, our redundant approach increases our confidence in our results revealing the Pir-to-ACx pathway, which are also consistent with some previous studies (Costa et al., 2017; Tasaka et al., 2020). Yet while our experiments yielded robust and reproducible results using these strategies, the question remains why some other research groups previously did not observe this projection. One explanation could simply relate to differences in the tracer used (retroAAVs here versus fluorescein-labeled dextran (Budinger et al., 2006; Budinger and Scheich, 2009)) and/or the animal model (mice here versus gerbils (Budinger et al., 2006; Budinger and Scheich, 2009)). In addition, due to the sparseness of the Pir-to-ACx projection, it is possible that this pathway may have gone unnoticed in some previous studies in which it was not the target of investigation.

Some previous rodent studies examined auditory-olfactory integration in ACx, providing an important foundation for understanding where and how the brain integrates auditory and olfactory stimuli (Cohen et al., 2011; Cohen and Mizrahi, 2015; Gilday and Mizrahi, 2023). Our data build upon this foundation and underscore modulation of sound responses by odors in naïve, passively exposed, awake mice. Previous experiments using more complex stimuli, such as modulation of ACx responses to pup vocalizations by pup odors, used acetophenone as a control odor stimulus, showing no effect on sound responses (Cohen et al., 2011). That said, the Cohen et al. study was performed in anesthetized animals, and it has since been well-established that neuronal responses vary greatly in anesthetized versus awake states (Fontanini and Katz, 2008; Lesicko et al., 2022). Our results suggest that olfactory modulation by simple odorants such as acetophenone may depend on brain state. Furthermore, the combined olfactory and auditory stimuli in previous studies were not bound precisely in time, and often involved other factors such as behavioral training and associative learning that could have altered cross-modal coding (Cohen et al., 2011; Cohen and Mizrahi, 2015; Gilday and Mizrahi, 2023). Our results in passively exposed, untrained mice suggest that auditory-olfactory integration in ACx does not require behavioral training. Future studies will be needed to further disentangle the complex relationships between brain state, stimulus complexity, and experience-dependent learning involved in auditory-olfactory integration and its neurophysiological underpinnings. Interestingly, an important recent study has suggested that auditory-olfactory integration occurs in higher-order areas of entorhinal cortex due to shared auditory and olfactory inputs (Wu et al., 2023). Our results add to this growing body of literature and suggest that auditory-olfactory integration occurs at even earlier stages in the sensory processing hierarchy than previously realized.

In sound-responsive odor-modulated units, we found that odor alone produces a slight enhancement of firing rate even in the absence of sound (Figs. 4K, 5I, 6F). Thus, olfactory input may affect both baseline and sound-evoked ACx activity. Our findings of both enhanced and suppressed sound-evoked responses are consistent with previous studies in other modalities such as in audio-visual integration, showing that visual stimuli evoke both excitation and modulation of ACx (Bizley et al., 2007; Kayser et al., 2008; Atilgan et al., 2018). However, the relatively modest degree of the odor-mediated effects on ACx activity observed here (Figs. 3-6), coupled with the sparse nature of the Pir-to-ACx projection (Figs. 1-2), overall suggest that this olfactory input to ACx likely sculpts auditory processing in ACx, rather than providing a strong excitatory drive.

We found that Pir input to ACx contributes to odor-mediated enhancement of baseline and sound-evoked responses, but not suppression of sound-evoked responses. It is intriguing to consider what mechanism might account for this dichotomy. Because most long-range intra-telencephalic projections are excitatory (Shepherd, 2013; Harris and Shepherd, 2015), odor enhancement may be due to activation of ACx by direct Pir input, whereas odor suppression may arise from indirect input via local circuitry or via other brain regions. In our anterograde tracing experiments identifying Pir-recipient neurons, we observed that the TeA receives a large amount of Pir input (Fig. 2), a finding that is consistent with previous studies (Tasaka et al., 2020, 2023) and suggests that TeA is a region that might relay Pir input to ACx, ultimately affecting ACx activity. Together these results pave the way for future studies to disentangle the relative contributions of direct Pir input versus indirect pathways to auditory-olfactory integration in ACx, and how the Pir-to-ACx pathway interacts with other multimodal and higher-order brain circuity.

Finally, it is worth noting that previous studies of auditory-olfactory integration have largely focused on parental behavior, including the effects of pup odors on auditory processing in maternal mice, as well as pup retrieval (Cohen and Mizrahi, 2015; Marlin et al., 2015; Marlin and Froemke, 2017; McRae et al., 2023). Moving forward, it will be interesting to determine the extent to which the Pir-to-ACx pathway may contribute to parental behavior, pup retrieval, and other complex behaviors. Although our present study has focused on the effects of simple neutral odorants, future studies are poised to investigate the additional effects of ethologically relevant odors, such as pup odors, social odors, food odors, or predator odors. Furthermore, auditory-olfactory integration in ACx is known to undergo plasticity during motherhood as well as during formation of odor-sound associations (Cohen and Mizrahi, 2015; Gnaedinger et al., 2019; Gilday and Mizrahi, 2023). Drawing from these findings, we suggest that the Pir-to-ACx pathway serves as a novel contributor to multimodal processing that may play a role in multisensory plasticity and learning.

